# SwiftTCR: Efficient computational docking protocol of TCRpMHC-I complexes using restricted rotation matrices

**DOI:** 10.1101/2024.05.27.596020

**Authors:** Farzaneh M. Parizi, Yannick J. M. Aarts, Nils Smit, Dona Roran AR, Daniëlle Diepenbroek, Wieke A. Krösschell, Levin Thijs, Joost Tepperik, Sanna Eerden, Dario F. Marzella, Gayatri Ramakrishnan, Li C. Xue

## Abstract

The T cell’s ability to discern self and non-self depends on its T cell receptor (TCR), which recognizes peptides presented by MHC molecules. Understanding this TCR-peptide-MHC (TCRpMHC) interaction is important for cancer immunotherapy design, tissue transplantation, pathogen identification, and autoimmune disease treatments. Understanding the intricacies of TCR recognition, encapsulated in TCRpMHC structures, remains challenging due to the immense diversity of TCRs (>10^8^/individual), rendering experimental determination and general-purpose computational docking impractical. Addressing this gap, we have developed a rapid integrative modeling protocol leveraging unique docking patterns in TCRpMHC complexes. Built upon PIPER, our pipeline significantly cuts down FFT rotation sets, exploiting the consistent polarized docking angle of TCRs at pMHC. Additionally, our ultra-fast structure superimposition tool, GradPose, accelerates clustering. It models a case in 3-4 minutes on 12 CPUs, showcasing a speedup of up to 25-40 times compared to the ClusPro webserver. On a benchmark set of 38 TCRpMHC class I (TCRpMHC-I) complexes, our protocol outperforms the state-of-the-art docking tools in model quality. This protocol can potentially provide structural information to TCR repertoires targeting specific peptides. Its computational efficiency can also enrich existing pMHC-specific single-cell sequencing TCR data, facilitating the development of structure-based deep learning (DL) algorithms. These insights are essential for understanding T cell recognition and specificity, advancing the development of therapeutic interventions.

## 1. Introduction

T cells discern between self and non-self antigens through their T cell receptors’ (TCRs) recognition of peptide antigens presented by major histocompatibility complexes (MHCs). Endogenous peptides expressed by MHC class I (MHC-I) can activate CD8+ cytotoxic T cells, and exogenous antigens expressed by MHC class II (MHC-II) can activate CD4+ T cells. Understanding the interaction between TCR-peptide-MHC (TCRpMHC) has broad applications in immunology, autoimmune diseases (Zhong et al., 2013), infectious disease treatment (Dolton et al., 2022) and cancer immunotherapy(Waldman et al., 2020).

The fundamental question of how T cells recognize antigens within a variety of peptide antigens with high specificity has been the subject of considerable research effort (Jerne, 1971; Scott-Browne et al., 2009; Van Laethem et al., 2012). This inquiry has revealed an evolving understanding that T cell activation relies on the simultaneous recognition of foreign peptide antigens and self MHC molecules, with a focus on reconciling selection and germline-encoded models (Zhong et al., 2013). Structural investigations into T cell surveillance have yielded valuable insights into this inquiry. Notably, all (with a few exceptions (Gras et al., 2016)) existing experimentally determined TCRpMHC structures demonstrate that TCRs maintain consistent polarity over pMHC (**Fig. 1**), which influences TCR specificity and binding affinity, and is also shown to impact T cell activation and signaling in mice and humans (Zareie et al., 2021). For instance, TCRs with reversed polarity in a viral epitope-specific T cell repertoire were unable to support TCR signaling due to lymphocyte-specific protein tyrosine kinase (Lck) mislocalization in the presence of the CD8+ coreceptor (Zareie et al., 2021). Structural investigation has also contributed to tumor-associated or tumor-specific peptide target identification (Antunes et al., 2019) and the optimization of cancer immunotherapies (Saotome et al., 2023), which holds the promise to reduce the toxicity of TCR therapies caused by TCR cross-reactivity to self-peptides.

**Figure 1.**
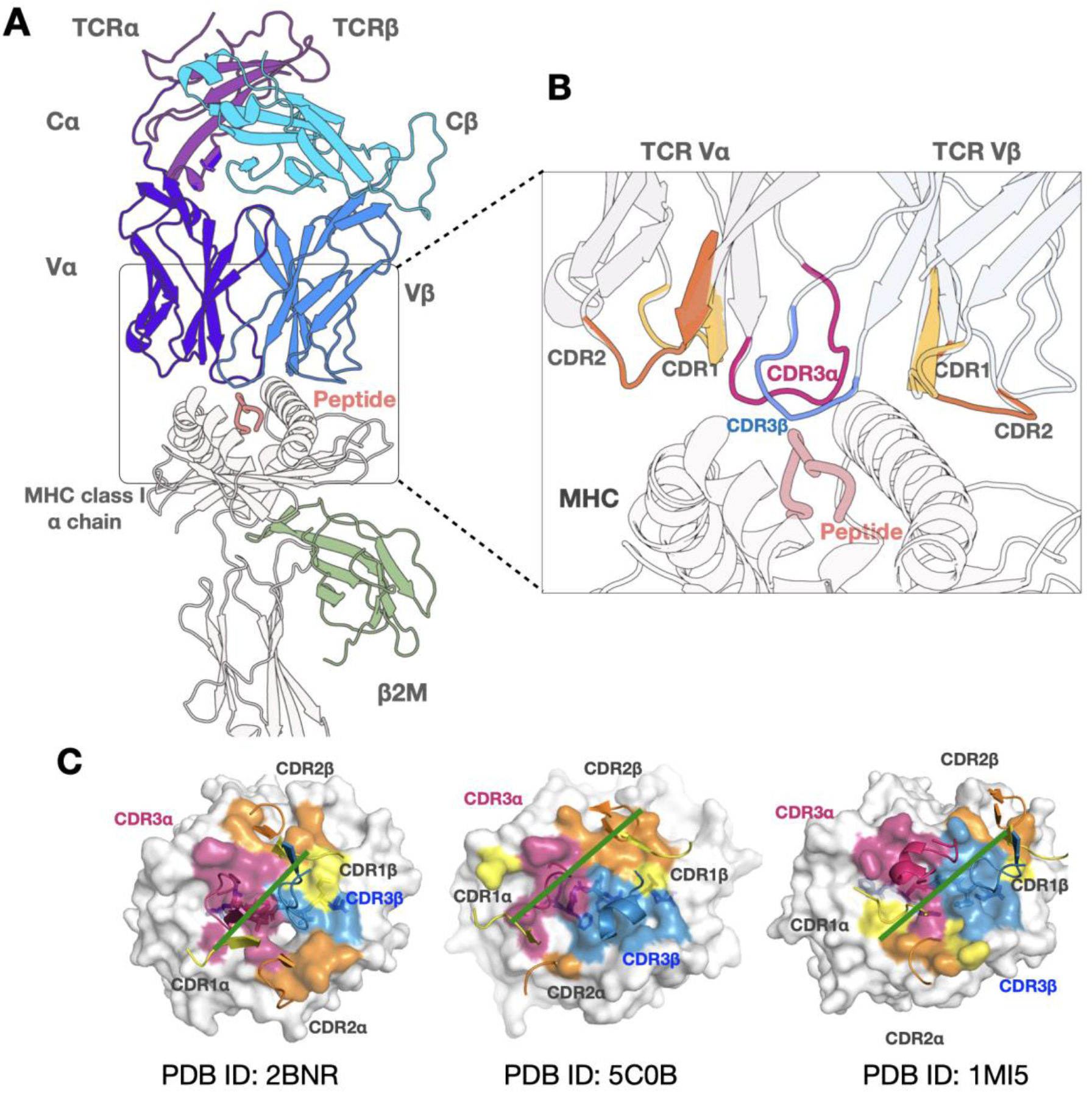
Structural visualization of TCR binding to pMHC. **A**) A TCRpMHC structure. The peptide (pink) binds inside the binding groove of the MHC molecule (white) and is presented on the cell surface to be visible to T cells (shades of blue and purple). The MHC-I molecule consists of two domains: alpha chain and beta-2 microglobulin (β2m) (green). The TCR, comprising alpha and beta chains with constant domains (Cα, purple; Cβ, cyan) binding to the T cell’s surface and variable domains (Vα, dark purple; Vβ, blue), interacting with the pMHC. The experimental structure (PDB ID: 2BNR) is illustrated in the figure, visualized with Protein Imager (Tomasello et al., 2020). **B**) The TCR has three CDR loops, CDR1 and CDR2 typically interact with the MHC’s conserved alpha helices to stabilize the interaction, CDR3 plays a major role by recognizing the peptide antigen presented by the MHC as self or non-self. Binding interface of TCRpMHC complex, where TCR Vα (light purple) positions over MHC-I α2 helix (white) with its CDR3α (magenta) above the N-terminal half of the peptide (salmon). TCR Vβ positions over MHC-I α1 helix, with its CDR3β (blue) extending over the C-terminal portion of the peptide. Images made by Protein Imager (Tomasello et al., 2020) for an experimental structure (PDB ID: 2BNR). **C)** The colors indicate the docking footprints of the TCR CDR loops (CDR1: yellow, CDR2: orange, CDR3α: magenta, CDR3β: skyblue) on their cognate pMHC complex surface (white). The green line on the surface shows the docking angle, representing the polarity of TCR over pMHC for three different experimental structures.

The TCR structure is a heterodimer composed of α and β chains (or γ and δ chains in γδ T cells). Each chain consists of a variable (V) domain and a constant (C) domain (**Fig. 1A**). The V domains participate in antigen recognition, whereas the C domains assist in the TCR’s anchoring to the T cell’s surface. Each variable domain of a TCR contains three complementarity-determining region (CDR) loops (CDR1-3). The TCR heterodimer docks diagonally with the long axis of the MHC peptide-binding groove. CDR1-2 loops primarily interact with MHC, whereas CDR3 interacts with the peptide, contributing uniquely to TCR specificity (**Fig. 1B, C**). Structures provide atomistic details of the TCRpMHC interaction, as well as insights into interaction energies and physicochemical properties. With increasing advances in artificial intelligence (AI) approaches, structure-based TCR specificity predictions may hold the promise to decipher the long-elusive TCR recognition code. The integration of structural information into AI methods has been shown to enhance the development of generalizable predictive approaches for MHC-binding peptide predictions and drug design (Marzella et al., 2023). However, the available experimental TCRpMHC structures (∼350 experimental structures in PDB (Berman, 2000)/ IMGT (Kaas et al., 2004)) are not enough to train a powerful AI predictor, necessitating the development of computational modeling methods that encompass the immense diversity of TCR, MHC, and peptide sequences.

A few computational tools have been developed for 3D modeling of TCRpMHC complexes, based on docking, AI, or homology modeling. Docking predicts complex structures from individual protein structures by generating tens of thousands of docking conformations and selecting the ones with lowest energies (the hallmark of native conformations). Flexible docking allows conformational changes for potentially better model quality but requires much more computational time, while rigid body docking maintains fixed conformations for faster modeling. A recent benchmark study (Peacock & Chain, 2021) assessed the performance of four docking platforms —ClusPro (Kozakov et al., 2017), ZDOCK (Pierce et al., 2014), and HADDOCK (de Vries et al., 2010)— in modeling TCRpMHC interactions. This study, utilizing an expanded benchmark set of 44 TCRpMHC docking cases, revealed that HADDOCK demonstrated superior performance. Further, ClusPro performed effectively in terms of model quality and computing time. ClusPro uses PIPER (Kozakov et al., 2006) for sampling and a closed source clustering method. AI-based tools like Immunebuilder (Abanades et al., 2023) and TCRmodel2 (Yin et al., 2023) predict TCR structures, with TCRmodel2 additionally capable of modeling entire TCRpMHC complexes. Another AI-based docking pipeline (Bradley, 2023) uses hybrid structural templates for sampling docking modes, similar to homology modeling. Homology modeling tools, exemplified by PANDORA (Parizi et al., 2023) and APE-Gen (Fasoulis et al., 2024), generate pMHC complex structures and LYRA (Klausen et al., 2015) for modeling TCR structures. Additionally, ImmuneScape (Li et al., 2019) is designed to model the entire TCRpMHC structure.

While AI-based tools have potential, their performance on novel TCRpMHCs remains uncertain. Nonetheless, standalone TCR and pMHC modeling tools show promise (Abanades et al., 2023; Fasoulis et al., 2024; Parizi et al., 2023; Yin et al., 2023). However, for TCRpMHC modeling, by leveraging the wealth of information known for TCR binding to pMHC, it is possible to incorporate these into 3D modeling for more efficient and accurate modeling.

Exploiting the docking polarity of TCRs over pMHC (**Fig. 1C**), we developed SwiftTCR, an integrative framework built on top of the PIPER software by only sampling on the feasible conformational space (**Fig. 2**). Tailored for TCRpMHC modeling, our framework reduces rotation sets of Fast Fourier Transform (FFT) from 202,491 to 3,775 rotations, aligning with the canonical binding orientation of TCR and pMHCs. This allows more in-depth sampling in the feasible conformational space with improved modeling results. With several other innovations (see Methods/Results), our protocol is up to 25-40 times faster than ClusPro, and is suitable for large-scale TCRpMHC modeling. Benchmarking on 38 experimentally resolved TCRpMHC-I complexes revealed that our method competes with state-of-the-art docking tools in terms of model quality and speed. With this acceleration, we envision our software delivering large-scale 3D models of TCRpMHC conformations closely aligned with experimental structure orientations, facilitating the design of more generalizable deep learning (DL) algorithms for TCR specificity predictions, unrestricted by species or allele types.

**Figure 2.**
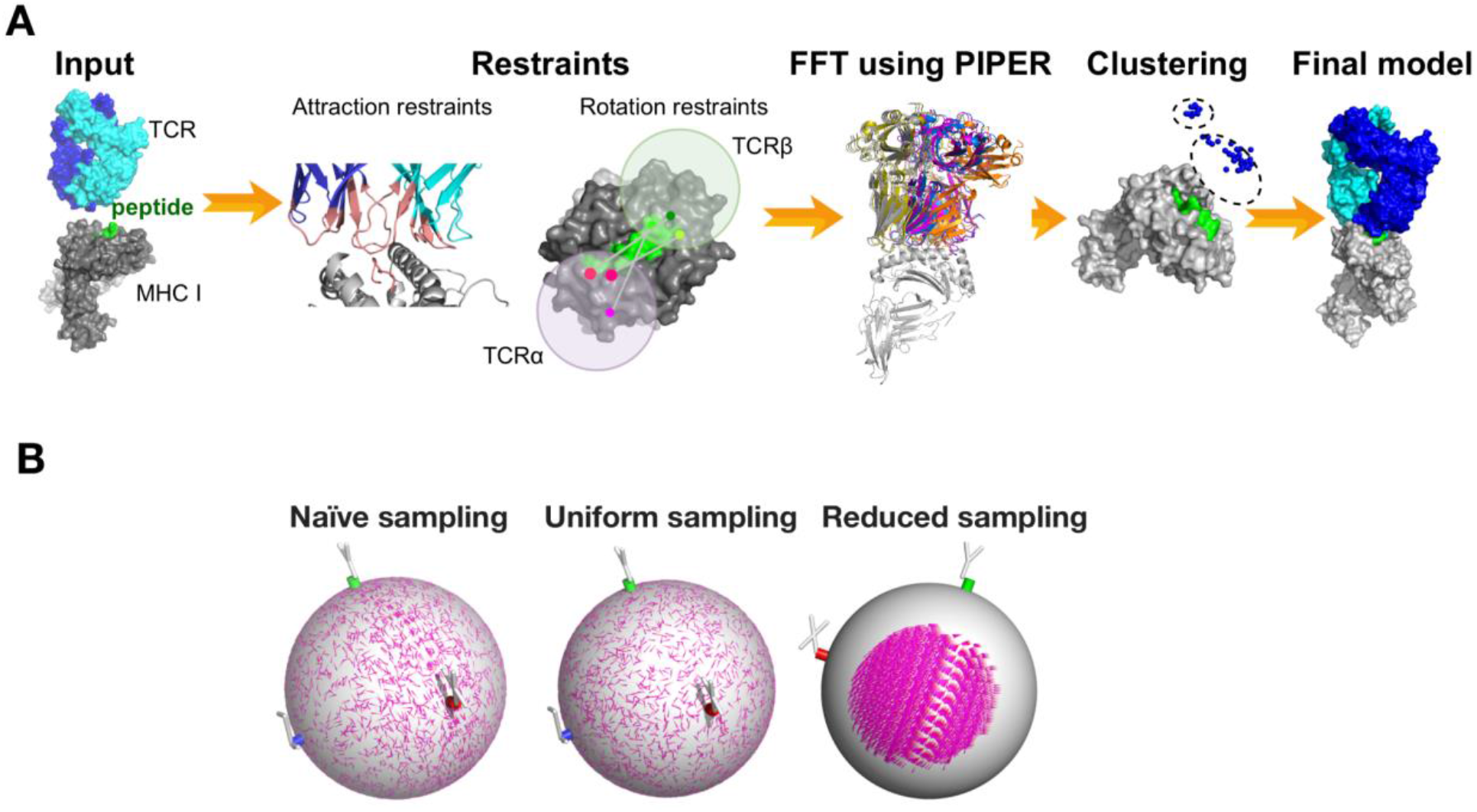
Schematic overview of SwiftTCR docking pipeline for modeling TCRpMHC complexes. **A)** Given the conformation of TCR and pMHC structures, the pipeline generates TCRpMHC structures. Initial orientation of pMHC and TCR to a reference plane, followed by docking using PIPER with a customized rotation set for consistent TCR docking angles to pMHC and specified attraction to peptide and CDR3 residues. Subsequently, clustering of the models is performed using our fast pairwise i-RMSD calculation, and the top-ranked models are selected to identify near-native complexes. **B)** Sampled rotations on a 3D sphere. Naïve sampling: is generated by taking even step sizes on the three Euler angles, leading to uneven distribution (more dense sampling along the X-Y plane) and oversight of critical regions. Uniform Rotation Sampling using quaternions: rotation matrices are generated uniformly across a 3D sphere. Reduced sampling: rotation matrices generated in B are filtered based on crossing angles and incident angles observed in the experimental TCRpMHC structures. For easy visualization, only 5000 random rotation matrices are shown for naïve sampling and uniform sampling. The reduced sampling contains a total of 3,775 rotations, all of which are shown here.

## 2. Methods

We established a pipeline for TCRpMHC modeling using PIPER docking software. In our docking approach, we imposed two key restrictions: 1) limiting the buried surface area to TCR CDR loops and peptide regions, and 2) constraining rotational space to ensure consistent polarized docking angles of TCRs at pMHC. This enhanced both the speed and accuracy of the docking process. In clustering, we leverage GradPose (Rademaker et al., 2023), our efficient structure superimposition tool. **Fig. 2A** depicts the general pipeline architecture used in this work.

### 2.1 Benchmark Dataset

To evaluate SwiftTCR, we used a TCR benchmark set (Peacock & Chain, 2021) which includes 44 cases of experimental TCRpMHC class I and II complexes, where for each of the bound TCRpMHC structures the corresponding unbound TCR and unbound pMHC structures were available. The structures were IMGT (Kaas et al., 2004) numbered. In this proof-of-concept study, we have restricted our analyses to MHC class I alone, which amounts to a total of 38 cases of Human (36) and mouse (2) alleles. The docking cases were labeled as Easy, Medium, assessing interface variability between bound and unbound experimental structures (see **Supplementary Method 1.3**).

### 2.2 SwiftTCR docking procedure

#### 2.2.1 Docking prerequisite

SwiftTCR can take input PDB files with any chain IDs and any numbering as it automatically renumbers the TCRs using ANARCI (Dunbar & Deane, 2016) and renames the chain IDs for TCR, peptide, MHC automatically. The input TCR must have complete variable domains, as the variable domains were used to calculate the crossing angle and incident angles for the reduced rotation matrices.

#### 2.2.2 Defining crossing angle and incident angle

Taking into account the TCR docking polarity, our algorithm conducts sampling within the favorable binding angle ranges, for speed and better model quality. The binding angle was characterized by two parameters: the crossing angle and the incident angle (Rudolph et al., 2006) (**Fig. 3**).

**Figure 3.**
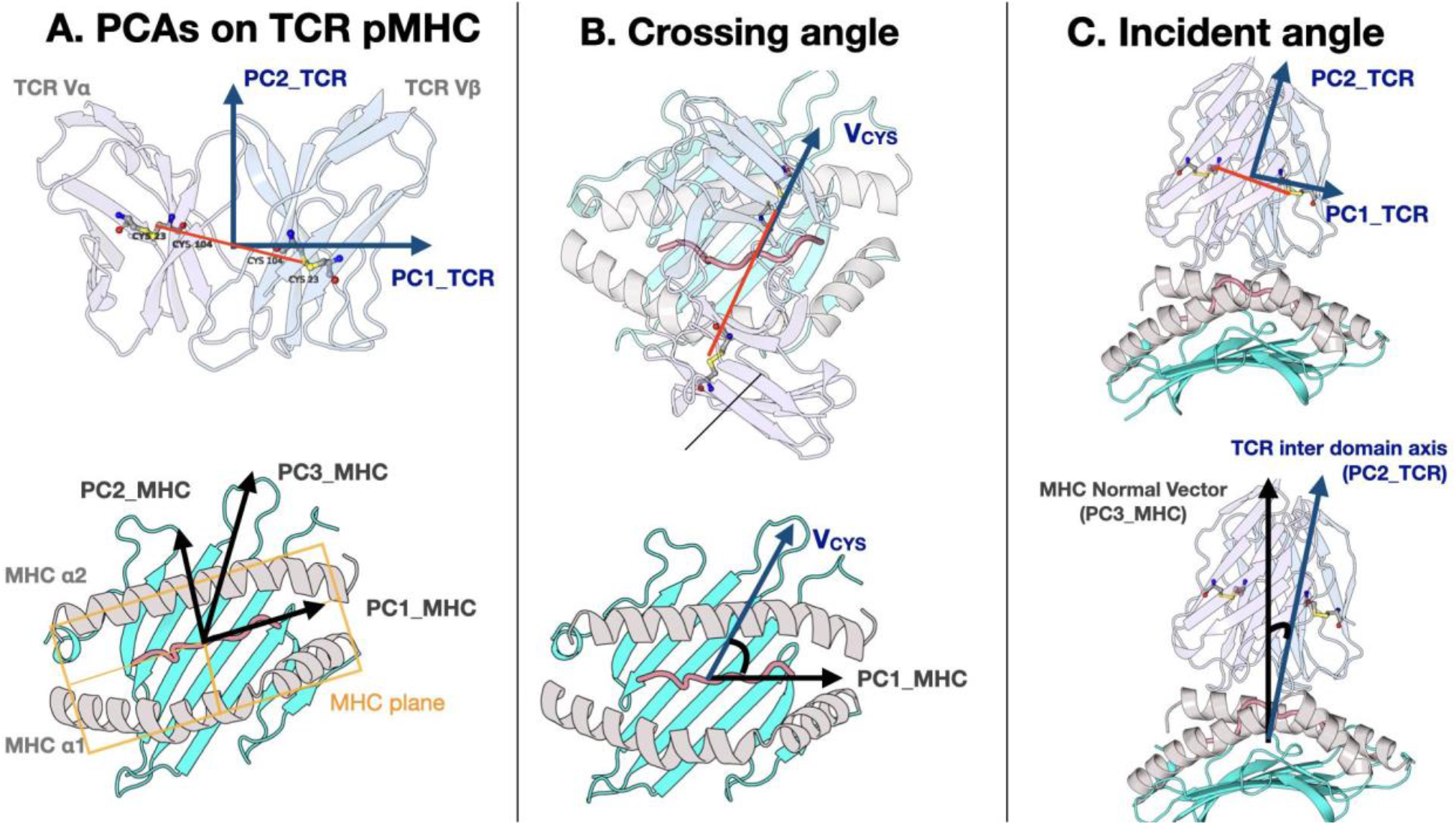
The definition of crossing angle and incident angle. **A**) PCA applied on TCR variable domain. To take into account the TCR docking polarity, PC1_TCR always points from TCR Vα to TCR Vβ. PC2_TCR: TCR interdomain axis of rotation points from TCR variable domain to TCR constant domain. More defined, V_CYS: the TCR docking vector which is the centroid of the four conserved cysteines of TCR variable domains (shown by the red lines crossing these residues) also points from TCR Vα to TCR Vβ. The MHC plane is defined by PCA analysis on the MHC helices. PC1_MHC: The long axis of the MHC binding groove, pointing from the N-ter of the peptide to the C-ter of the peptide. PC2_MHC: The width axis of the MHC binding groove, pointing to the MHC α1 helix. PC1_MHC and PC2_MHC define the MHC plane. PC3_MHC: The norm vector of the MHC plane, pointing to TCR. **B**) The definition of crossing angle, which measures the angle between the TCR docking vector (V_CYS) and the long axis of the MHC binding groove (PC1_MHC). Thus, a reversed polarity will have a negative crossing angle. **C**) The definition of incident angle, which measures the tiltness of TCR over the MHC plane. Incident angle is the angle between the TCR interdomain axis of rotation (PC2_TCR) and the MHC normal vector (PC3_MHC).

The crossing angle measures the docking polarity or the twist of the TCR over the pMHC, which is the angle between the TCR docking vector (centroid of conserved disulfide bonds) and the vector of the MHC binding groove (**Fig. 3B**). Specifically, we applied principal component analysis (PCA) on the MHC α1 and α2 helices (IMGT number: 50-86 and 140-176) to derive the first principal component (PC1_MHC, pointing from N-ter of the peptide to the C-ter) as the binding groove vector (**Fig. 3A**). To calculate the TCR docking vector, we took four conserved cysteines of the TCR (IMGT number: 23, 104 for both chains), and use the vector pointing from the cysteine geometric center on chain D to the one on chain E as the TCR docking vector (V_CYS). The angle between PC1_MHC and V_CYS is the crossing angle (see **Supplementary Fig. S1**).

The incident angle measures the tilt of the TCR variable domain over the pMHC, which is the angle between the normal vector of the MHC peptide binding groove plane (PC3_MHC, pointing outside the MHC and towards the TCR) and the TCR interdomain rotation axis (PC2_TCR, pointing from the TCR variable domains to the constant domains) (**Fig. 3C**). To get the TCR interdomain rotation axis, we conducted Principal Component Analysis (PCA) on the variable domain of TCR, and PC2_TCR which is the TCR interdomain rotation axis (**Fig. 3A**). Our tool uses only the variable region of TCR, making it compatible with single-cell sequencing studies that might lack constant region data.

### 2.3. Definition of restraints

#### 2.3.1 Defining reduced docking rotation matrices for favorable docking angles

Uniform sampling is essential for our docking method to ensure unbiased exploration of docking orientations in 3D space. To achieve this, we generated uniformly distributed rotations using Shoemake’s group theory-based algorithm (Shoemake, 1992) (**Supplementary Algorithm S1, Fig. 2B**). We retained rotation matrices that produced favorable docking angles for TCRpMHC complexes: specifically, crossing angles within [15°, 90°] and incident angles within [0°, 35°], based on data from 177 TCRpMHC crystal structures (**Supplementary Fig. S1**). This filtering yielded 3,775 valid rotation matrices. The final matrices were saved as a rotation file (.prm) for input to PIPER, which then applies FFT to identify optimal translations for the given rotations. For details, see **Supplementary Method 1.1**.

#### 2.3.2. Defining attractive restraints for binding residues

We further enhance the docking accuracy by restricting the conformational search space with additional attraction terms targeting the defined interface residues (**Fig. 4A**). During docking calculations, an additional attractive force is applied to the selected residues. These constraints guide the docking process towards selecting the appropriate pose. We added attraction to all peptide residues and all CDR residues (**Fig. 4A**). The objective is to ensure that during sampling, conformations where the peptide is not in close proximity to TCRs are assigned a relatively unfavorable energy score.

**Figure 4.**
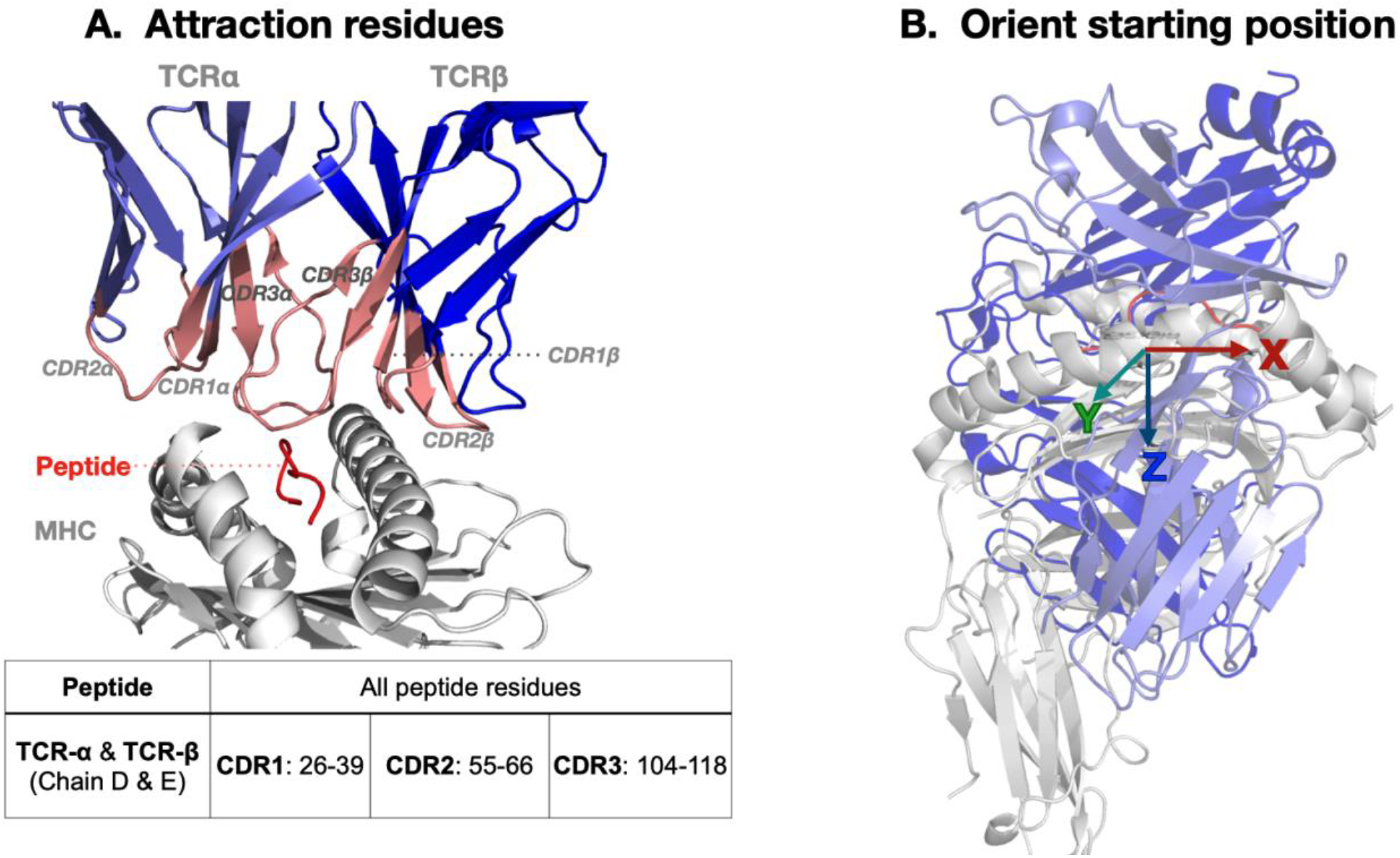
Preparatory insights to SwiftTCR. **A)** Attractive Residues highlighted by pink marking key residues in the TCR (purple) binding interface to the peptide (red) bound to MHC (white), guiding docking around this interface. Attractive Residues Table lists peptide residues and corresponding TCR CDR1-3 residues for binding, following IMGT numbering. **B)** Initial docking position illustrates pMHC and TCR structures’ initial orientation, centered on their principal components at the Cartesian coordinate origin, exemplified by PDB ID 1AO7.

We add these attraction residues by modifying the PDB files to include Attraction in column 55-60 of ATOM records of PDB files (0= no attraction, 1= attraction), which were then used as input for PIPER.

### 2.4. Docking

#### 2.4.1 Preparation for electrostatic calculations

We used PDB2PQR (Dolinsky et al., 2007) for the preparation of input PDB files. This tool converts PDB files of biomolecular structures into PQR files, and adds hydrogens to optimize the hydrogen bonding network.

#### 2.4.2. Alignment to Reference Structure

Before docking, all TCR and pMHC structures were aligned to the reference complex 2BNR using PyMOL (Schrödinger, LLC, 2015). Alignment was based on superposed Cα atoms to ensure chain correspondence and consistent orientation. This standardization enabled accurate application of the reduced rotation matrices. An example starting position is shown in **Fig. 4B**.

#### 2.4.3 Docking with reduced rotation matrices and attractive restraints

Docking was performed in PIPER, treating the TCR as the ligand and the pMHC as the receptor. The search used the reduced rotational space (**Fig. 2A**) and defined attractive restraints between peptide and CDR3 residues (**Fig. 4A**). From 3,775 sampled orientations, the top 1,000 conformations were selected based on energy for clustering.

#### 2.4.5 Clustering

Docked complexes were clustered using a greedy algorithm based on pairwise interface RMSD (i-RMSD), similar to ClusPro (Kozakov et al., 2017) but with differing thresholds. Interface residues were defined as TCR atoms within 10 Å of any pMHC atom. Specifically, we calculate all pairwise i-RMSD for all top 1000 low-energy models. Similar to ClusPro, greedy clustering is employed to rank conformations based on their adherence to the specified threshold. Clustering initiated from the lowest-energy model; all models within a 3 Å cutoff were grouped and excluded from subsequent rounds. The process continued until all models were assigned or the cluster number limit (100) was reached. Clusters were ranked by size, with ties resolved by lowest energy.

For efficiency, GradPose (Rademaker et al., 2023) was integrated to massively accelerate i-RMSD calculations and structural alignments.

### 2.5 Evaluations

Model quality was assessed using Critical Assessment of PRedicted Interactions (CAPRI) metrics: Ligand RMSD (L-RMSD), interface RMSD (i-RMSD), and fraction of native contacts (f_nat_) (see **Supplementary Methods 1.5**). Models were classified into *High, Medium, Acceptable*, or *Incorrect* quality categories (**Supplementary Table S2**).

## 3. Results

### 3.1 SwiftTCR generates good-quality models in minutes on CPUs

To evaluate performance, we tested SwiftTCR on the TCRpMHC benchmark dataset, which includes 38 experimentally determined TCR–pMHC-I structures (Peacock & Chain, 2021). Docked conformations were compared against their corresponding experimental structures. The same benchmark was used in the latest evaluation of TCRpMHC docking tools, which assessed: ClusPro(Kozakov et al., 2017), LightDock (Jiménez-García et al., 2018), ZDOCK (Pierce et al., 2014), and HADDOCK (de Vries et al., 2010). Among these, ClusPro showed the most consistent performance, employing PIPER, an FFT-based docking algorithm combined with clustering to identify native-like interactions. As SwiftTCR also uses PIPER with a reduced rotational sampling strategy for efficient TCRpMHC docking, we selected ClusPro as the primary method for comparison.

The success plot for generating acceptable, medium-, and high-quality models among the top-ranked predictions are shown in **Fig. 5A**. While ClusPro achieved competitive performance in previous benchmarks, SwiftTCR outperformed it under default settings (clustering cutoff: ClusPro 9 Å; SwiftTCR 3 Å). Because SwiftTCR uses a reduced rotational sampling space, a smaller clustering cutoff was required. Benchmark tests (data not shown) revealed that a 3 Å cutoff yielded higher-quality models, especially within the top 10 ranks, by increasing cluster diversity and improving model selection.

**Figure 5.**
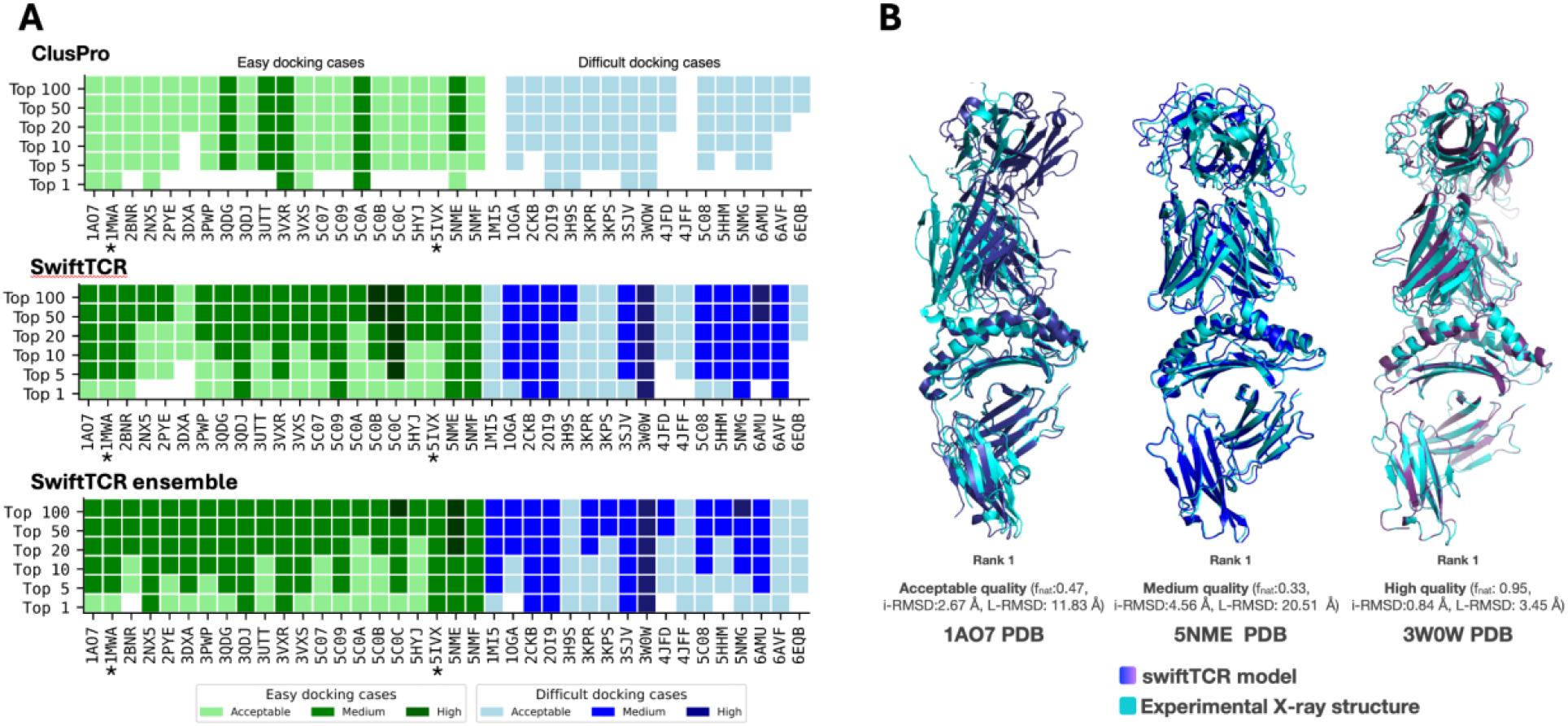
The unbound-unbound docking performance of SwiftTCR in modeling TCRpMHC-I interactions on a benchmark set with 38 experimental TCRpMHC-I structures (Peacock & Chain, 2021). **A)** Comparison with ClusPro in terms of success plot. Top panel: ClusPro performance, taken from (Peacock & Chain, 2021). Middle panel: SwiftTCR performance using unbound x-ray structures as input. Bottom panel: SwiftTCR using ensemble of 5 TCR models generated by AlphaFold3 and unbound x-ray pMHC structure as input. A 3Å clustering threshold is used for SwiftTCR. Model quality was assessed based on L-RMSD, i-RMSD, and f_nat_ metrics, following CAPRI criteria. Cases indicated by * are mouse alleles. **B)** Example of a 3D model generated by SwiftTCR shown in shades of purple and blue, compared to experimental structures in cyan.

SwiftTCR performed robustly, producing at least one acceptable-quality model among the top 10 ranked predictions in 97.4% (37 out of 38) benchmark cases (**Fig. 5A middle panel**). Within the top 50 models, over 81% of cases included at least one medium- or high-quality result. Visualization of representative models across quality levels (acceptable, medium, high) shows that the predicted TCRpMHC binding modes closely align with experimental structures (**Fig. 5B**).

### 3.2 Ensemble input conformations improve SwiftTCR’s performance

A key limitation of SwiftTCR is that it performs rigid-body docking and does not account for the intrinsic flexibility of the CDR loops. This is evident in several challenging cases (e.g., 1MI5, 3KPR, 4JFD, 4JFF, and 6EQB), where few medium-quality models are found among the top 1,000 predictions (**Supplementary Fig. S2**). Comparison of bound and unbound structures shows substantial conformational changes in CDR2 or CDR3 upon binding (**Supplementary Fig. S3**), highlighting the importance of loop flexibility.

To address this, we generated an ensemble of five TCR conformations (for details, see **Supplementary Method 1.2**) and used them, together with the unbound X-ray pMHC structure, as input for SwiftTCR. Docking models from the five runs were pooled, clustered, and the top 100 models selected for evaluation. This ensemble strategy improved performance in most cases, increasing the success rate of medium-quality models in the top 20 predictions from 71.1% to 81.6% (**Fig. 5A**, bottom panel). Notably, SwiftTCR-ensemble obtained 100% success rate in top 5 predictions.

**Figure 6** illustrates the details of the ensemble docking for the PDB structure 1MI5. While docking with the unbound X-ray TCR structure failed to produce medium-quality models, the AF-predicted TCR structures yielded substantially more medium-quality solutions. Structural comparison suggests that this difference arises from steric incompatibilities: the unbound X-ray TCR conformation exhibits clashes with the pMHC, whereas the CDR3 loops in the AF-predicted models adopt conformations that avoid these clashes. This improved conformational compatibility likely facilitates the generation of higher-quality docking solutions with correct global orientations, even though AF3 does not fully capture the native bound conformations of the CDR3 loops.

### 3.2 Accelerating TCRpMHC modeling: SwiftTCR as the fastest physics-based method

The computational time of FFT-based docking approaches largely depends on the number of rotation matrices used. By default, PIPER employs 70,000 rotations, taking 27 minutes on a server with 30 CPU cores to dock one case without pairwise distance restraints. Because we focus on sampling around the valid interfaces, we are able to do much more dense sampling (step size of 6 degrees), resulting in 202,491 unique rotations for the 3D sphere, which is reduced to 3,775 rotations after applying the docking angle and incident angle filters. This allows SwiftTCR to generate models for one TCRpMHC modeling case in 3 to 4 minutes with better quality on 12 CPUs (11th Gen Intel(R) Core(TM) i7-11800H @ 2.30GHz) locally.

**Figure 6.**
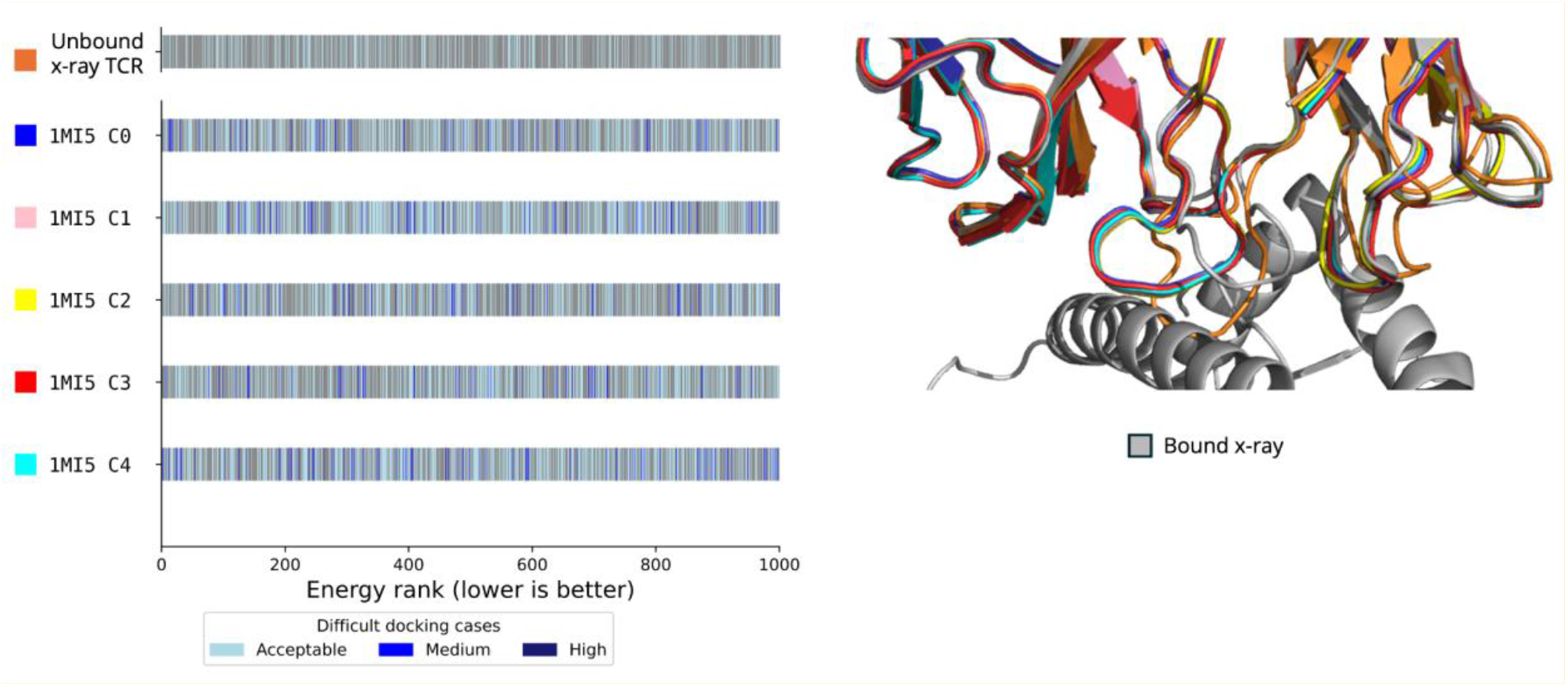
Ensemble docking on PDB entry 1MI5. Five TCR models (1MI5_cX, X = 1–5) were generated using AF3, and each combined with the unbound X-ray pMHC structure was used as input for SwiftTCR, resulting in five independent docking TCRpMHC runs. Left: The top 1,000 models per run are displayed as Melquiplots, where each line represents a model ranked by SwiftTCR energy and colored by quality (darker = better; gray = incorrect). For comparison, the top 1,000 models from a docking run using the unbound X-ray TCR structure are also shown. Right: Superposition of the input TCR structures with the bound conformation highlights conformational changes. The unbound TCR conformation (orange) is shown clashing with the pMHC.

State-of-the-art docking tools, such as the HADDOCK and ClusPro servers, offer user-friendly interfaces but often require several hours to complete the docking process for a single TCRpMHC case. In contrast, ZDock, another FFT-based docking tool, operates significantly faster, completing each case in 5–15 minutes on their web server. However, ZDock employs a 15-degree sampling step, which is considerably less dense than SwiftTCR’s 6-degree sampling. This lower sampling density is likely a factor in ZDock’s suboptimal performance in TCRpMHC docking. The speed and accuracy of SwiftTCR makes large-scale modeling of TCRpMHC interactions feasible, especially for AI-driven applications such as structural TCR specificity prediction.

### 3.3 Comparison with AlphaFold3

To enable an objective comparison with AlphaFold3 (AF3) (Abramson et al., 2024), we assembled a benchmark dataset comprising 17 PDB structures released after AF3’s training cutoff date (September 30, 2021). The dataset includes structures containing TCR variable genes TRAV and TRBV, together with MHC genes, as well as TCR gene combinations that were not present in the PDB prior to the AF3 training date (**Supplementary Table S1**).

Because SwiftTCR requires structural inputs, we first used AF3 to generate five TCR models per case and employed PANDORA to generate one pMHC structure, which together served as input for SwiftTCR (see details in **Supplementary Materials 1.2 and 1.3**).

AF3 demonstrates robust performance on this dataset, producing at least one medium-quality model among its top five predictions in 70.6% of cases and at least an acceptable-quality model in 100% of cases (**Fig. 7**). SwiftTCR complements AF3 by substantially increasing the success rate of medium-quality models—from 70.6% within AF3’s top five predictions to 94.1% within SwiftTCR’s top 100 models. However, the results also indicate that reliably ranking medium-quality models within the top 1–5 predictions remains challenging for SwiftTCR.

**Figure 7.**
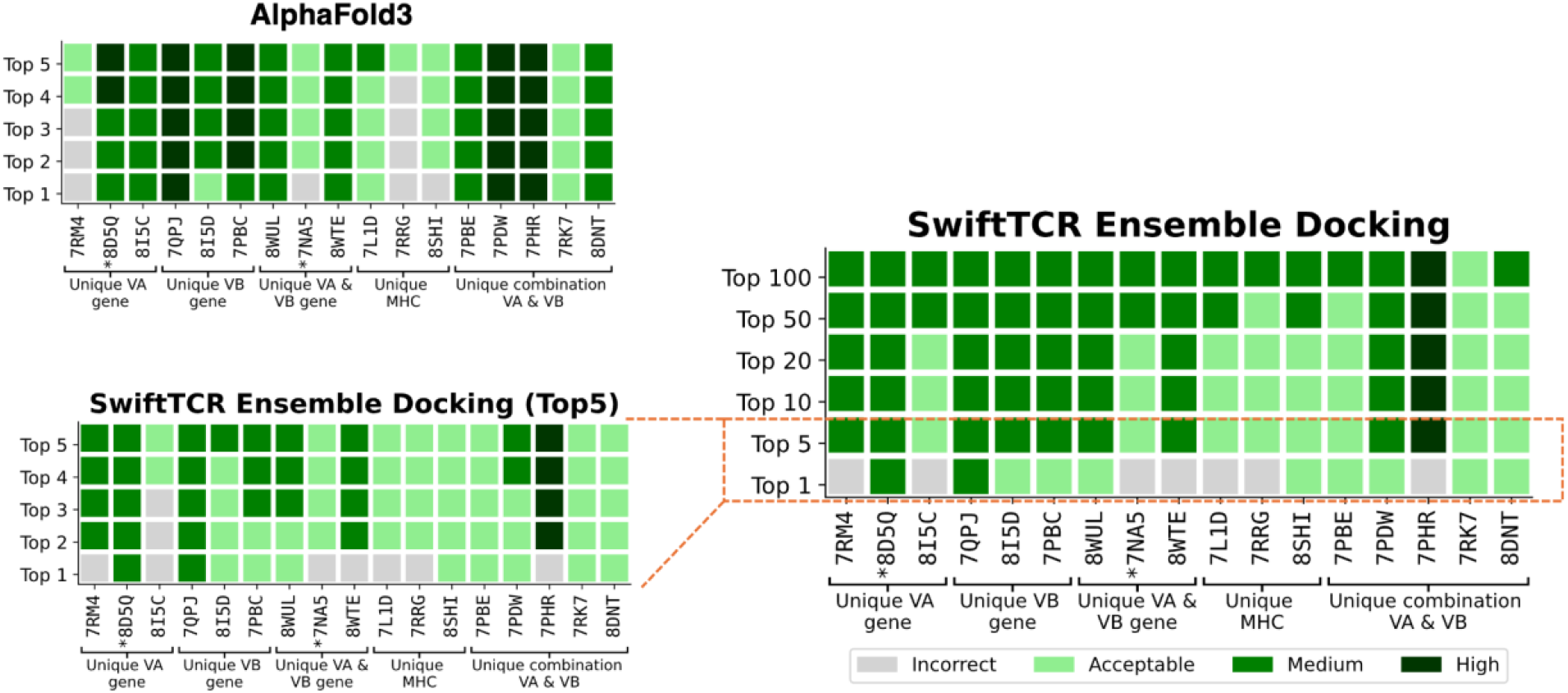
The comparison with AlphaFold3 on 17 cases with unique V-gene or MHC. SwiftTCR complements AF3 by improving the success rate of medium-quality models—from 70.6% among AF3’s 5 models to 94.1% among SwiftTCR’s top 100 models. At the same time, it highlights that selecting medium-quality models within the top 1–5 remains challenging for SwiftTCR. For each case, SwiftTCR was executed five times using five distinct TCR structures and one pMHC structural input, generating a total of 5,000 models, from which the top 100 are reported; and AF3 was run once, performing five independent diffusion-based structure generation processes and producing five models in total.

## 4. Discussion

Three-dimensional TCRpMHC structures are invaluable for deciphering the long-elusive TCR recognition code. To this end, we have developed SwiftTCR, a fast and reliable integrative rigid-body docking software for modeling TCRpMHC complexes. Its unique features include uniform sampling for evenly distributed docking orientations, reduced rotation sets for TCRpMHC interactions, and a modular Python design. Our protocol achieves rapid modeling speeds (∼3 to 4 min on 12 CPUs), outperforming state-of-the-art physics-based docking tools in model quality and computational efficiency. Using ensemble inputs to capture TCR flexibility further enhances SwiftTCR’s performance (up to 94% success rate for medium-quality models in top 100 predictions), inspired by Giulini et al.’s work on improving antibody–antigen docking (Xu et al., 2024).While this study currently focuses on MHC class I, it can be readily expanded to MHC-II by specifying rotation sets suitable for TCRpMHC-II structures, a direction currently under investigation.

SwiftTCR is built upon PIPER (Kozakov et al., 2006) for sampling and energy calculation for TCRpMHC modeling. SwiftTCR utilizes a dual criterion for selecting top models: prioritizing energy first (preferring lower values), followed by cluster sizes (preferring larger clusters, and reporting the cluster center as the representative model) to efficiently group similar models, aiding in the selection of near-native TCRpMHC orientations. The idea of using cluster size to rank the models was originally used in ClusPro (Kozakov et al., 2017). The rationale is that docking typically samples more models in the lowest energy funnel.

While SwiftTCR is able to generate medium to high-quality models for most cases (82-94% cases), they are not necessarily ranked top (**Fig. 5A** and **Supplementary Fig. S3)**, indicating the need for improving our scoring scheme. Implementing a multi-step clustering approach, starting with large cutoffs to select the top 1-2 clusters and refining with smaller cutoffs, may also help. We evaluated this multi-step clustering approach in the recent CAPRI competition (Round 56), the target of which is a TCRpMHC complex. Our multi-step clustering approach achieved the second highest scoring (3/10 Medium quality, 5/10 acceptable quality models). Additionally, including coevolution information between TCR and MHC might help us to further narrow down the search space and improve both the sampling step, enabling the generation of more medium-high quality models, and the scoring step.

SwiftTCR complements AI-based modeling approaches such as AlphaFold and its variants (Abramson et al., 2024; Bradley, 2023; Jumper et al., 2021; Wu et al., 2025; Yin et al., 2023), which have achieved impressive results in modeling structures of single-chain proteins and protein complexes. Unlike these data-driven, statistics-based methods that rely on large-scale training data, SwiftTCR employs a physics-based docking framework and does not require prior training. As all statistical learning approaches, AlphaFold models may be susceptible to biases toward structural templates, alleles, or gene combinations overrepresented in the PDB databank – an effect illustrated, for example, in comparisons between AlphaFold2 and the physics-based method PANDORA (Marzella et al., 2022). In contrast, physics-based approaches are guided by energy functions rather than learned statistical patterns, which may enhance their generalizability to novel TCRs, peptides, and MHC alleles, as reflected by SwiftTCR’s high success rate in generating medium-quality models within its top 100 predictions (**Fig. 7**). In addition, AlphaFold3 is computationally demanding: when provided with sequence input, the default implementation requires approximately 1 hour and 50 minutes on a single A100 GPU, while an accelerated version still requires roughly 4 minutes per case to generate five models. By comparison, given input TCR and pMHC structures, SwiftTCR generates 1,000 docked models in approximately 3–4 minutes on 12 CPUs. Moreover, its FFT-based sampling procedure is inherently parallelizable and could, in principle, be substantially accelerated on GPUs, which are well suited for large-scale parallel FFT computations.

It is important to note that SwiftTCR is designed to predict the 3D structures of TCRpMHC complexes given a known interacting pair, rather than to predict binding specificity or discriminate true binders from non-binders. While docking scores are reported, they should not be interpreted as measures of binding likelihood. Assessing the model on mismatched pairs is therefore beyond the scope of this study. Extending SwiftTCR to evaluate specificity and reduce false positives is an important direction for future work.

### Limitations

In our analysis, we observed that longer CDR3 loops (≥10 amino acids) yielded less acceptable quality models in the top 1000 sampled models (**Supplementary Fig. S2**). In addition, CDR2 conformations also greatly impact the docking performance (PDB ID: 2NX5, 4JFD, 6EQB, **Supplementary Fig. S3**). Using an ensemble of input TCR structures has helped to improve the docking performances (**Fig. 5 and 6**).

Given SwiftTCR’s rigid-body docking nature, post-docking flexible refinement is required to capture binding-induced conformational changes of TCR loops upon pMHC recognition. While incorporating TCR ensembles as input partially accounts for receptor flexibility and substantially improves performance (**Fig. 5A**), a light, localized refinement step (e.g., short loop relaxation in solvent) is expected to further enhance model quality. In future work, we will evaluate the impact of such refinement using HADDOCK3 (Giulini, Reys, et al., 2025).

We note that SwiftTCR requires individual structures of TCRs and pMHCs as input, which can currently be obtained using tools such as AlphaFold (Abramson et al., 2024), TCRmodel2 (Yin et al., 2023) or MODELLER (Webb & Sali, 2016) for TCRs and PANDORA (Marzella et al., 2022) for pMHCs. Generating these structures can be time-consuming. To address this, we are developing ultra-fast AI tools capable of modeling these structures in milliseconds. For instance, our task-specific model SwiftMHC (Baakman et al., 2025) predicts all-atom pMHC structures and binding affinities in just 0.009 s per case for HLA-A*02:01 9-mer peptides when run in batch mode. In addition, our diffusion-based model, MHC-Diff (Frühbuß et al., 2025), accurately models pMHC structures (currently, α-carbon only) while capturing peptide flexibility. We are actively extending these approaches to support additional MHC alleles and TCRs.

## Data and Code availability

We will make the data available on Zenodo with DOIs upon the acceptance of the manuscript. And the SwiftTCR code is freely available on GitHub: https://github.com/X-lab-3D/swifttcr.

### Abbreviations

AI: Artificial Intelligence
DL: Deep Learning
FFT: Fast Fourier Transform
PCA: Principal Component Analysis
CAPRI: Critical Assessment of PRedicted Interactions
3D: Three-Dimensional
PDB: Protein Data Bank
MHC: Major Histocompatibility Complex
pMHC: Peptide-MHC
TCR: T Cell Receptor
TCRpMHC-I: TCR-pMHC class I
CDR: Complementarity-Determining Region
RMSD: Root Mean Square Deviation
i-RMSD: Interface Root Mean Square Deviation
L-RMSD: Ligand Root Mean Square Deviation
f_nat_: Fraction of native contacts

## Acknowledgment

FM is supported by the Kika grant (grant number 454). LX is supported by Hanarth Fond. GR was supported by Europees Fonds voor Regionale Ontwikkeling (EFRO) (R0005582). The computation resources are funded by SurfSara (grant numbers EINF2380). We would like to express our gratitude to the ClusPro team and George Jones at Boston University and Stony Brook University for generously sharing the PIPER code and their help, which facilitated the implementation of restraints in our work. We also thank Kevin J van Geemen and Daniel T Rademaker, for their helpful discussions, for making the clustering step fast. We thank Prof. Alexandre Bonvin at Utrecht University for proposing the concept of ensemble docking. We are also grateful to E. Vermeulen for preparing the dataset used for comparison with AF3.

## Authors Contribution

Conceived the study: GR, YA, FP and LX. Project supervision and coordination: LX. Manuscript writing: FP and LX. Manuscript editing: YA, LX, GR, NS, SE, DD and DM. Software development: YA, DM and NS. Benchmark evaluation: YA, FP, NS, DR, JT, LT and SE. Ensemble evaluation: DD, WK, LT. Figures: FP, YA, NS, DR, DD, LT, WK and LX. All authors reviewed and approved the final manuscript.

All authors read and contributed to the manuscript. FM and YA agree that the order of their respective names may be changed for personal pursuits to best suit their own interests.

## Key Points

- Developed a high-speed, physics-based docking tool for TCRpMHC complexes, capable of completing runs in minutes on 12 CPUs (with potential for further acceleration on GPUs).
- Implemented a group theory-based algorithm to generate uniformly distributed rotation matrices, crucial for robust clustering-size-based scoring.
- Reduced the docking search space from 202,491 to 3,775 rotations by leveraging the docking polarity of TCRpMHC interactions.

## Biographical Note

Li C. Xue, PhD, is an Assistant Professor in Biomedical Sciences at Radboud University Medical Center. Her group develops AI methods for protein structure modeling in rational cancer immunotherapy design.

## References

Abanades, B., Wong, W. K., Boyles, F., Georges, G., Bujotzek, A., & Deane, C. M. (2023). ImmuneBuilder: Deep-Learning models for predicting the structures of immune proteins. Communications Biology, 6(1), Article 1.10.1038/s42003-023-04927-7

Abramson, J., Adler, J., Dunger, J., Evans, R., Green, T., Pritzel, A., Ronneberger, O., Willmore, L., Ballard, A. J., Bambrick, J., Bodenstein, S. W., Evans, D. A., Hung, C.-C., O’Neill, M., Reiman, D., Tunyasuvunakool, K., Wu, Z., Žemgulytė, A., Arvaniti, E., … Jumper, J. M. (2024). Accurate structure prediction of biomolecular interactions with AlphaFold 3. Nature, 630(8016), 493–500. 10.1038/s41586-024-07487-w

Antunes, D. A., Abella, J. R., Devaurs, D., Rigo, M. M., & Kavraki, L. E. (2019). Structure-based Methods for Binding Mode and Binding Affinity Prediction for Peptide-MHC Complexes. Current Topics in Medicinal Chemistry, 18(26), 2239–2255. 10.2174/1568026619666181224101744

Baakman, C., Crocioni, G., Geng, C., Rademaker, D. T., Frühbuß, D., Aarts, Y. J. M., & Xue, L. C. (2025). SwiftMHC: A High-Speed Attention Network for MHC-Bound Peptide Identification and 3D Modeling. Immunology. 10.1101/2025.01.20.633893

Berman, H. M. (2000). The Protein Data Bank. Nucleic Acids Research, 28(1), 235–242. 10.1093/nar/28.1.235

Bradley, P. (2023). Structure-based prediction of T cell receptor:peptide-MHC interactions. eLife, 12, e82813. 10.7554/eLife.82813

de Vries, S. J., van Dijk, M., & Bonvin, A. M. J. J. (2010). The HADDOCK web server for data-driven biomolecular docking. Nature Protocols, 5(5), Article 5. 10.1038/nprot.2010.32

Dolinsky, T. J., Czodrowski, P., Li, H., Nielsen, J. E., Jensen, J. H., Klebe, G., & Baker, N. A. (2007). PDB2PQR: Expanding and upgrading automated preparation of biomolecular structures for molecular simulations. Nucleic Acids Research, 35(Web Server), W522–W525. 10.1093/nar/gkm276

Dolton, G., Rius, C., Hasan, M. S., Wall, A., Szomolay, B., Behiry, E., Whalley, T., Southgate, J., Fuller, A., Morin, T., Topley, K., Tan, L. R., Goulder, P. J. R., Spiller, O. B., Rizkallah, P. J., Jones, L. C., Connor, T. R., & Sewell, A. K. (2022). Emergence of immune escape at dominant SARS-CoV-2 killer T cell epitope. Cell, 185(16), 2936-2951.e19. 10.1016/j.cell.2022.07.002

Dunbar, J., & Deane, C. M. (2016). ANARCI: Antigen receptor numbering and receptor classification. Bioinformatics, 32(2), 298–300. 10.1093/bioinformatics/btv552

Fasoulis, R., Rigo, M. M., Lizée, G., Antunes, D. A., & Kavraki, L. E. (2024). APE-Gen2.0: Expanding Rapid Class I Peptide–Major Histocompatibility Complex Modeling to Post-Translational Modifications and Noncanonical Peptide Geometries. Journal of Chemical Information and Modeling, 64(5), 1730–1750. 10.1021/acs.jcim.3c01667

Frühbuß, D., Baakman, C., Teusink, S., Bekkers, E., Jegelka, S., & Xue, L. C. (2025). Fast and Accurate Peptide–MHC Structure Prediction via an Equivariant Diffusion Model. Bioinformatics. 10.1101/2025.04.28.650973

Giulini, M., Reys, V., Teixeira, J. M. C., Jiménez-García, B., V. Honorato, R., Kravchenko, A., Xu, X., Versini, R., Engel, A., Verhoeven, S., & Bonvin, A. M. J. J. (2025). HADDOCK3: A Modular and Versatile Platform for Integrative Modeling of Biomolecular Complexes. Journal of Chemical Information and Modeling, 65(13), 7315– 7324. 10.1021/acs.jcim.5c00969

Giulini, M., Xu, X., & Bonvin, A. M. (2025). Improved structural modelling of antibodies and their complexes with clustered diffusion ensembles. Bioinformatics. 10.1101/2025.02.24.639865

Gras, S., Chadderton, J., Del Campo, C. M., Farenc, C., Wiede, F., Josephs, T. M., Sng, X. Y. X., Mirams, M., Watson, K. A., Tiganis, T., Quinn, K. M., Rossjohn, J., & La Gruta, N. L. (2016). Reversed T Cell Receptor Docking on a Major Histocompatibility Class I Complex Limits Involvement in the Immune Response. Immunity, 45(4), 749–760. 10.1016/j.immuni.2016.09.007

Jerne, N. K. (1971). The somatic generation of immune recognition. European Journal of Immunology, 1(1), 1–9. 10.1002/eji.1830010102

Jiménez-García, B., Roel-Touris, J., Romero-Durana, M., Vidal, M., Jiménez-González, D., & Fernández-Recio, J. (2018). LightDock: A new multi-scale approach to protein–protein docking. Bioinformatics, 34(1), 49–55. 10.1093/bioinformatics/btx555

Jumper, J., Evans, R., Pritzel, A., Green, T., Figurnov, M., Ronneberger, O., Tunyasuvunakool, K., Bates, R., Žídek, A., Potapenko, A., Bridgland, A., Meyer, C., Kohl, S. A. A., Ballard, A. J., Cowie, A., Romera-Paredes, B., Nikolov, S., Jain, R., Adler, J., … Hassabis, D. (2021). Highly accurate protein structure prediction with AlphaFold. Nature, 596(7873), 583–589. 10.1038/s41586-021-03819-2

Kaas, Q., Ruiz, M., & Lefranc, M. (2004). IMGT/3Dstructure-DB and IMGT/StructuralQuery, a database and a tool for immunoglobulin, T cell receptor and MHC structural data. Nucleic Acids Research, 32(Suppl_1), D208–D210. 10.1093/nar/gkh042

Klausen, M. S., Anderson, M. V., Jespersen, M. C., Nielsen, M., & Marcatili, P. (2015). LYRA, a webserver for lymphocyte receptor structural modeling. Nucleic Acids Research, 43(W1), W349–W355. 10.1093/nar/gkv535

Kozakov, D., Brenke, R., Comeau, S. R., & Vajda, S. (2006). PIPER: An FFT-based protein docking program with pairwise potentials. Proteins: Structure, Function, and Bioinformatics, 65(2), 392–406. 10.1002/prot.21117

Kozakov, D., Hall, D. R., Xia, B., Porter, K. A., Padhorny, D., Yueh, C., Beglov, D., & Vajda, S. (2017). The ClusPro web server for protein–protein docking. Nature Protocols, 12(2), 255–278. 10.1038/nprot.2016.169

Li, S., Wilamowski, J., Teraguchi, S., Van Eerden, F. J., Rozewicki, J., Davila, A., Xu, Z., Katoh, K., & Standley, D. M. (2019). Structural Modeling of Lymphocyte Receptors and Their Antigens. In S. Kaneko (Ed.), In Vitro Differentiation of T-Cells (Vol. 2048, pp. 207–229). Springer New York. 10.1007/978-1-4939-9728-2_17

Marzella, D. F., Crocioni, G., Radusinovic, T., Lepikhov, D., Severin, H., Bodor, D. L., Rademaker, D. T., Lin, C., Georgievska, S., Kessler, A. L., Lopez-Tarifa, P., Buschow, S., Bekkers, E., & Xue, L. C. (2023). Improving generalizability for MHC-binding peptide predictions through structure-based geometric deep learning (p. 2023.12.04.569776). bioRxiv. 10.1101/2023.12.04.569776

Marzella, D. F., Parizi, F. M., Tilborg, D. V., Renaud, N., Sybrandi, D., Buzatu, R., Rademaker, D. T., ‘T Hoen, P. A. C., & Xue, L. C. (2022). PANDORA: A Fast, Anchor-Restrained Modelling Protocol for Peptide: MHC Complexes. Frontiers in Immunology, 13, 878762.

Parizi, F. M., Marzella, D. F., Ramakrishnan, G., ‘t Hoen, P. A. C., Karimi-Jafari, M. H., & Xue, L. C. (2023). PANDORA v2.0: Benchmarking peptide-MHC II models and software improvements. Frontiers in Immunology, 14. https://www.frontiersin.org/articles/10.3389/fimmu.2023.1285899

Peacock, T., & Chain, B. (2021). Information-Driven Docking for TCR-pMHC Complex Prediction. Frontiers in Immunology, 12. https://www.frontiersin.org/articles/10.3389/fimmu.2021.686127

Pierce, B. G., Wiehe, K., Hwang, H., Kim, B.-H., Vreven, T., & Weng, Z. (2014). ZDOCK server: Interactive docking prediction of protein–protein complexes and symmetric multimers. Bioinformatics, 30(12), 1771–1773. 10.1093/bioinformatics/btu097

Rademaker, D. T., Van Geemen, K. J., & Xue, L. C. (2023). GradPose: A very fast and memory-efficient gradient descent-based tool for superimposing millions of protein structures from computational simulations. Bioinformatics, 39(8), btad444. 10.1093/bioinformatics/btad444

Rudolph, M. G., Stanfield, R. L., & Wilson, I. A. (2006). How Tcrs Bind Mhcs, Peptides, and Coreceptors. Annual Review of Immunology, 24(1), 419–466. 10.1146/annurev.immunol.23.021704.115658

Saotome, K., Dudgeon, D., Colotti, K., Moore, M. J., Jones, J., Zhou, Y., Rafique, A., Yancopoulos, G. D., Murphy, A. J., Lin, J. C., Olson, W. C., & Franklin, M. C. (2023). Structural analysis of cancer-relevant TCR-CD3 and peptide-MHC complexes by cryoEM. Nature Communications, 14(1), Article 1. 10.1038/s41467-023-37532-7

Scott-Browne, J. P., White, J., Kappler, J. W., Gapin, L., & Marrack, P. (2009). Germline-encoded amino acids in the αβ T-cell receptor control thymic selection. Nature, 458(7241), 1043–1046. 10.1038/nature07812

Tomasello, G., Armenia, I., & Molla, G. (2020). The Protein Imager: A full-featured online molecular viewer interface with server-side HQ-rendering capabilities. Bioinformatics, 36(9), 2909–2911. 10.1093/bioinformatics/btaa009

Van Laethem, F., Tikhonova, A. N., & Singer, A. (2012). MHC restriction is imposed on a diverse T cell receptor repertoire by CD4 and CD8 co-receptors during thymic selection. Trends in Immunology, 33(9), 437–441. 10.1016/j.it.2012.05.006

Waldman, A. D., Fritz, J. M., & Lenardo, M. J. (2020). A guide to cancer immunotherapy: From T cell basic science to clinical practice. Nature Reviews Immunology, 20(11), Article 11. 10.1038/s41577-020-0306-5

Webb, B., & Sali, A. (2016). Comparative Protein Structure Modeling Using MODELLER. Current Protocols in Bioinformatics, 54(1). 10.1002/cpbi.3

Wu, F., Zhao, Y., Xiao, Y., Qin, C., Wang, F., Wu, Z., Huang, L.-K., Liu, X., Song, J., He, B., Rossjohn, J., & Yao, J. (2025). Fast and accurate modeling of TCR-peptide-MHC complexes using tFold-TCR (p. 2025.01.12.632367). bioRxiv. 10.1101/2025.01.12.632367

Xu, X., Giulini, M., & Bonvin, A. M. J. J. (2024). Improved prediction of antibody and their complexes with clustered generative modelling ensembles. Bioinformatics Advances, 5(1), vbaf161. 10.1093/bioadv/vbaf161

Yin, R., Ribeiro-Filho, H. V., Lin, V., Gowthaman, R., Cheung, M., & Pierce, B. G. (2023). TCRmodel2: High-resolution modeling of T cell receptor recognition using deep learning. Nucleic Acids Research, 51(W1), W569–W576. 10.1093/nar/gkad356

Zareie, P., Szeto, C., Farenc, C., Gunasinghe, S. D., Kolawole, E. M., Nguyen, A., Blyth, C., Sng, X. Y. X., Li, J., Jones, C. M., Fulcher, A. J., Jacobs, J. R., Wei, Q., Wojciech, L., Petersen, J., Gascoigne, N. R. J., Evavold, B. D., Gaus, K., Gras, S., … La Gruta, N. L. (2021). Canonical T cell receptor docking on peptide–MHC is essential for T cell signaling. Science, 372(6546), eabe9124. 10.1126/science.abe9124

Zhong, S., Malecek, K., Johnson, L. A., Yu, Z., Vega-Saenz de Miera, E., Darvishian, F., McGary, K., Huang, K., Boyer, J., Corse, E., Shao, Y., Rosenberg, S. A., Restifo, N. P., Osman, I., & Krogsgaard, M. (2013). T-cell receptor affinity and avidity defines antitumor response and autoimmunity in T-cell immunotherapy. Proceedings of the National Academy of Sciences of the United States of America, 110(17), 6973–6978. 10.1073/pnas.1221609110

Giulini, Marco, Xiaotong Xu, and Alexandre MJJ Bonvin. (2025). Improved structural modelling of antibodies and their complexes with clustered diffusion ensembles. bioRxiv: 2025-02. 10.1101/2025.02.24.639865

