## Supplementary Material for "SwiftTCR: Efficient computational docking protocol of TCRpMHC-I complexes using restricted rotation matrices"

#### 1. Supplementary Method

##### 1.1 Generate uniformly distributed rotations

We first generated 202,491 rotation matrices uniformly sampled (using a step size of 6°) and then retained only those rotation matrices that can generate favorable docking angles for the TCRpMHC complex, i.e., $crossing\ angle \in [15°, 90°]$ and $incident\ angle \in [0°, 35°]$, resulting in 3,775 valid rotation matrices based on data from 177 crystal structures of TCRpMHC complexes (**Suppl. Fig. S1**).

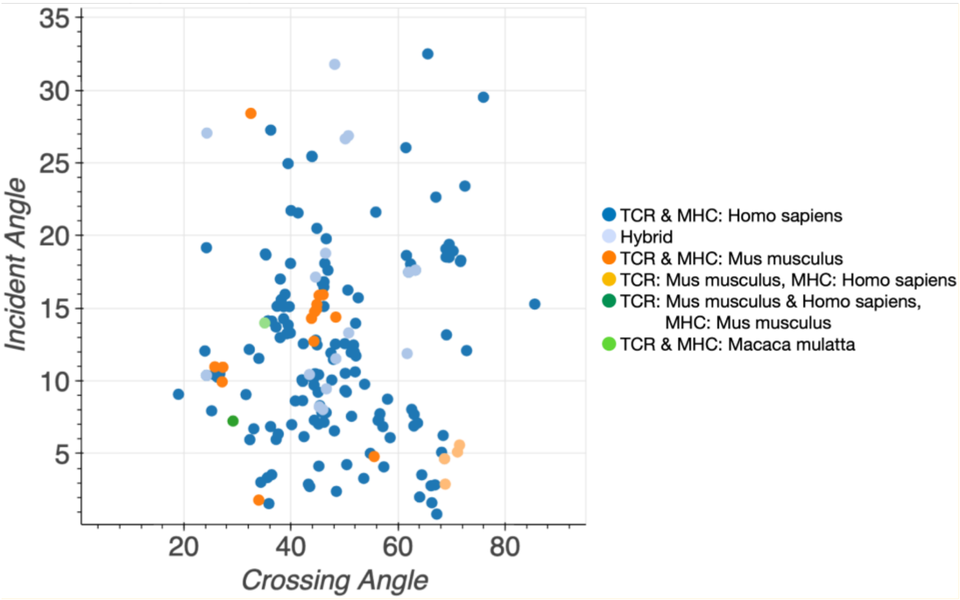

**Supplementary Figure S1. The incident and crossing angles observed in experimental TCRpMHC-I structures.** 177 experimental TCRpMHC-I structures in the IMGT/PDB (as of 06/06/2023) are retrieved from (Lin et al., 2025). Hybrid complexes are structures in which the TCR or MHC is composed of elements from multiple species. All the crossing angles fall within [15°, 90°], and the incident angles fall within [0°, 35°], demonstrating the docking polarity of TCRs. In SwiftTCR, we used these ranges for filtering the rotation matrices.

Generating uniformly distributed rotations is an important topic in robotics for sampling-based motion planning (Kuffner, 2004; Shoemake, 1992). Similarly, uniform sampling is crucial to our docking method to sample uniformly distributed docking orientations in 3D space for over-sampling or under sampling in certain areas to avoid biases. According to Euler's rotation theorem, any orientation can be described by three successive rotations about the three axes (x, y, z-axis). For example, rotating around the x-axis by $\theta$ degree, then around y-axis by $\phi$ degree and then around z-axis by $\eta$ degree. However, even sampling of three Euler angles $(\theta, \phi, \eta)$ result in unevenly distributed rotations (naïve sampling, **Fig. 2B**). We generated uniformly distributed rotations using the algorithm proposed by Shoemake based on group theory (Shoemake, 1992)(**Algorithm S1, Fig. 2B**). Uniform sampling ensures a uniform distribution across the entire space, enhancing precision, coverage and avoiding a bias towards specific orientations.

Once we generate rotation matrices that uniformly cover the 3D sphere, we filter these rotation matrices based on crossing angles and incident angles. For this we take a reference structure 2BNR to calculate docking angles after each rotation. We keep the pMHC structure fixed and use the generated rotation matrices to rotate the TCR and calculate the crossing angle and incident angle for the rotated TCRs. We used a range of [0, 35] degrees for the incident angle and a range of [15, 90] degrees for the crossing angle (these docking angle ranges are calculated on 177 experimental TCRpMHC structures in the PDB, **Fig. 5**). **Figure 2B** shows the reduced rotation set. This results in 3,775 rotation matrices that only include relevant rotations based on prior knowledge of existing TCRpMHC complexes in the PDB. The final rotation matrices are written as a rotation file (.prm), which is used as input for PIPER. PIPER uses FFT to find the best translation for the provided rotations.

---

**Algorithm S1:** uniformly distributed rotation matrices (quaternions)

---

**def** uniform_sampling (n_samples = 60):

1: $s = linspace(0{,}1, \text{n\_samples})$

2: $\sigma_1 = \sqrt{1-s}$

3: $\sigma_2 = \sqrt{s}$

4: $t_1 = linspace(0{,}1, \text{n\_samples})$

5: $t_2 = linspace(0{,}1, \text{n\_samples})$

6: $\theta_1 = 2\pi * t_1$

7: $\theta_2 = 2\pi * t_2$

### Define quaternion: [w, x, y, z]

$w = cos(\theta_2) * \sigma_2$
$x = sin(\theta_1) * \sigma_1$
$y = cos(\theta_1) * \sigma_1$
$z = sin(\theta_2) * \sigma_2$
**return** (w,x,y,z)

---

##### 1.2 SwiftTCR ensemble

Per TCR sequence pair 25 structures were predicted with AlphaFold3 (AF3) local installation using 5 random seed settings. The structures were converted to a pdb file and IMGT numbered with ANARCI (Dunbar & Deane, 2016). ProFit version 3.3 was used to calculate the pairwise RMSDs after aligning all the TCRs on their base structure excluding the CDR loops. The RMSD is used as a distance metric to perform agglomerative clustering of scikit-learn with complete linkage to generate five clusters. The cluster center model was selected as the representative model to dock with SwiftTCR. SwiftTCR is used to dock every combination of the selected 5 TCR models and the top pMHC model generated by PANDORA. These docking runs produce in total 5000 TCRpMHC structures, which are pooled and then clustered in SwiftTCR using the pairwise i-RMSD as distance metric, this time applied to the full set of 5,000 structures rather than the default 1,000. Finally, the cluster center models were selected for downstream evaluation.

##### 1.3 AlphaFold3

The performance of AF3was assessed using the local installation of AF3. The TCR and pMHC sequences were provided with a fixed seed of 1 and fixed chain IDs identical to the reference chain IDs. To evaluate the structures were the residue numbers of TCRβ shifted by 2000 residues and merged into chain D. MHC, beta-2 microglobulin (β2m) regions and the peptide structure were merged into chain A with MHC residue numbering shifted by 1000 and β2m by 2000 to preserve unique numbering and allow later lookup. These chainID naming and numbering is a requirement for SwiftTCRs evaluation code, which depends on DockQ (Basu & Wallner, 2016) to calculate the i-RMSD, L-RMSD and $f_{nat}$ to assess the final qualities.

##### 1.4 X-ray TCRpMHC structures with unique V-gene and MHC

To enable an objective comparison with AlphaFold3 (AF3), we assembled a benchmark dataset from TCR3d (Lin et al., 2025) comprising structures released after AF3's training cutoff date of September 30, 2021 (**Suppl. Table 1**). The benchmark includes 17 PDB structures containing TRAV, TRBV, and MHC genes, as well as TCR gene combinations not present in the PDB prior to the training cutoff and includes a large variation on crossing angles. All structures were standardized to ensure consistent chain annotation: using PyMOL (version 3.1) and pdb-tools (Rodrigues et al., 2018), chain IDs were harmonized such that MHC, β2m, and peptide were assigned to chains A, B, and C, respectively, and TCRα and TCRβ to chains D and E. TCR sequences were IMGT-numbered using ANARCI.

**Suppl. Table S1 The benchmark set used to compare AF3 and SwiftTCR.** It includes 17 PDB structures taken from the TCR3d database, containing TRAV, TRBV, and MHC genes, as well as TCR gene combinations not present in the PDB prior to the AF3 training cutoff of Sep. 30th 2021.

| PDB ID | MHC Name | Release date | TRAV gene | TRBV gene | Crossing angle (degrees) | Unique TRAV | Unique TRBV | Unique MHC |
|---|---|---|---|---|---|---|---|---|
| 7L1D | HLA-A*03 | 2022-03-23 | TRAV12-2 | TRBV9 | 47.4 | | | X |
| 7NA5 | H2-Db | 2022-06-22 | TRAV7-1 | TRBV2 | 57.2 | X | X | |
| 7PBC | HLA-A*02 | 2022-08-03 | TRAV12-2 | TRBV6-3 | 30.7 | | X | |
| 7PBE | HLA-A*02 | 2022-04-27 | TRAV12-1 | TRBV7-9 | 51.7 | | | |
| 7PDW | HLA-A*02 | 2022-08-03 | TRAV12-2 | TRBV6-2 | 33.2 | | | |
| 7PHR | HLA-A*02 | 2022-08-31 | TRAV17 | TRBV19 | 43.0 | | | |
| 7QPJ | HLA-A*02 | 2022-08-03 | TRAV12-2 | TRBV6-6 | 34.1 | | X | |
| 7RK7 | HLA-A*02 | 2022-11-02 | TRAV4 | TRBV10-3 | 80.7 | | | |
| 7RM4 | HLA-A*02 | 2022-02-09 | TRAV6 | TRBV11-2 | 35.3 | X | | |
| 7RRG | HLA-A*03 | 2022-03-23 | TRAV4 | TRBV11-2 | 73.8 | | | X |
| 8D5Q | H2-Ld | 2022-09-14 | TRAV6-7/DV9 | TRBV1 | 31.2 | X | | |
| 8DNT | HLA-A*02 | 2023-07-19 | TRAV12-2 | TRBV7-2 | 31.4 | | | |
| 8I5C | HLA-A*11 | 2023-08-23 | TRAV7-2 | TRBV13-2 | 47.7 | X | | |
| 8I5D | HLA-A*11 | 2023-08-23 | TRAV8-1 | TRBV16 | 44.5 | | X | |
| 8SHI | HLA-C*06 | 2023-06-28 | TRAV17 | TRBV6-5 | 21.4 | | | X |
| 8WTE | HLA-A*11 | 2024-05-01 | TRAV7-2 | TRBV2 | 64.1 | X | X | |
| 8WUL | HLA-A*11 | 2024-05-01 | TRAV7-2 | TRBV2 | 64.4 | X | X | |

#### 1.5 Evaluation metrics

##### Ligand RMSD (L-RMSD), i-RMSD and fraction of native contacts ($f_{nat}$)

We evaluated model qualities in terms of standard CAPRI metrics as L-RMSD, i-RMSD, and $f_{nat}$ using DockQ software (Basu & Wallner, 2016). Based on these evaluations, models were classified into High, Medium, Acceptable and Incorrect quality groups (**Suppl. Table 1**). RMSD values were computed based on backbone atoms (Cα, N, C, O), and interface residues were defined as those within a 10 Å distance of atoms on the other molecule.

For the L-RMSD calculations the TCR is considered as the receptor and the pMHC as the ligand. TCRs (both constant and variable domains) of the modeled TCRpMHC complex and the experimentally determined complex were superimposed to compute the RMSD between their pMHC conformations. Similarly, i-RMSD was calculated by superimposing the interfaces of the modeled TCRpMHC complex and the experimentally determined complex, followed by assessing the deviation of interface residues. $f_{nat}$ evaluates contact similarity between the modeled TCRpMHC and the experimental complex.

**Supplementary Table S2. Model qualification used based on the CAPRI** (Méndez et al., 2003) **scoring metrics.** Model qualities were categorized into high, medium, and acceptable quality groups according to CAPRI criteria for protein-protein docking quality evaluation (Lensink & Wodak, 2010).

| **Class** | **$F_{nat}$** | **L-RMSD (Å)** | **i-RMSD (Å)** |
|---|---|---|---|
| **High** | ≥ 0.5 and | "≤ 1.0 | Or ≤ 1.0" |
| **Medium** | ≥ 0.3 and | "≤ 5.0 | Or ≤ 2.0" |
| **Acceptable** | ≥ 0.1 and | "≤ 10.0 | Or ≤ 4.0" |
| **Incorrect** | ≤ 0.1 | - | - |

**Success Rate**

Following CAPRI (Lensink & Wodak, 2010), the model qualities are categorized based on how closely they resemble the experimental structure, classified as incorrect, acceptable, medium, or high-quality. The Success Rate is determined by calculating the percentage of cases where at least one successful model appears among the top M ranked models:

$$success\ rate\ (M) = \frac{num\ of\ successful\ cases\ when\ top\ M\ models\ are\ evaluated}{N}$$

Where N is the total number of cases (N = 38 in this study), and a case is counted as a success if at least one hit model (satisfying defined quality criteria (Lensink & Wodak, 2010)) appears among the top M ranked models. The quality criteria include high, medium, and acceptable quality models.

**1.6 Docking Difficulty Classification**

We follow the docking difficulty classification used in Peacock & Chain (2021), which employs standard interface-focused metrics. Cases are further described by flexibility: Easy cases (referred to as "rigid") have *i-RMSD ≤ 1.5 Å* and $f_{non\text{-}nat} \leq 0.4$, corresponding to complexes where the unbound (apo) structures undergo minimal conformational changes upon binding. Difficult cases (referred to as "medium" in Peacock & Chain (2021)) have either *i-RMSD > 1.5 Å and < 2.2 Å*, or *i-RMSD* > *1.5 Å* with $f_{non\text{-}nat}$ > *0.4*, representing complexes where the unbound (apo) structures undergo larger conformational changes, making docking more challenging.

#### 2. Supplementary Results

##### 2.1 Docking performance of top 1000 models

To assess whether lower-quality models in certain docking cases are due to sampling limitations or due to scoring, we examined the Melquiplot for the top 1,000 sampled models in each modeling case (**Suppl. Fig. S1**).

In this figure, each column shows 3D models for one docking case, with one line for one model. For each case, we generated up to 1,000 models, which are ranked by PIPER energy function (**Suppl. Fig. S1**). A perfect energy function will rank all good models to the top (top 1 is the best at the bottom of the bar). Good models are colored with green and blue, and wrong models are colored gray. With this plot, we can easily see how many good models are generated by sampling. We expect a higher presence of green models in each bar at the bottom, as top-ranked models generally exhibit higher quality.

Overall, SwiftTCR successfully generates a substantial number of medium to high-quality models in most cases. However, in certain instances, such as 2NX5, 1MI5, 3KPR, 3KPS, 4JFD and 4JFF, 6EQB suboptimal sampling was observed. We notice that the poor sampling of such cases can be explained by (**Suppl. Fig. S2**) (see details in the section below).

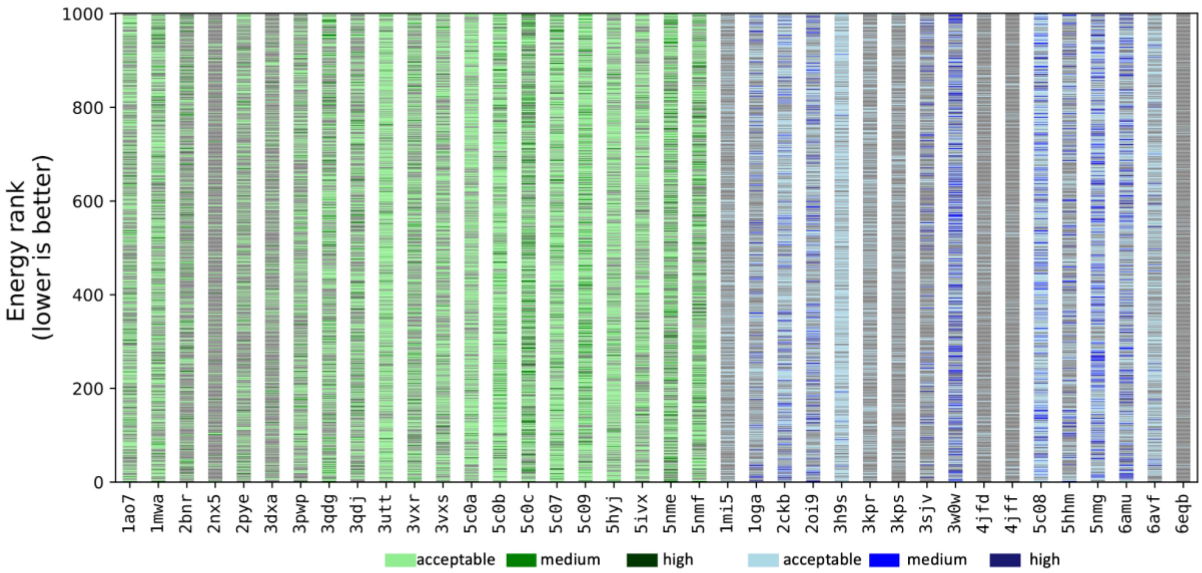


**Supplementary Figure S2: PIPER energy ranking of  the benchmark cases.** Shows the sampling performance across various modeling cases. Models are ranked by the PIPER energy function. The top 1000 models are selected, with the highest-ranked models at the bottom (lower PIPER energy). Each generated  model is color-coded in terms of standard CAPRI metrics by quality: light green/blue for acceptable quality, green/blue for medium quality, and dark green/blue for high quality.

#### 2.2 Long CDR3 loops or "wrong" CDR2 loop conformations may cause SwiftTCR to generate fewer good-quality models

To further evaluate the cases with poor sampling, we visually inspected the models to determine whether CDR3 lengths or the CDR loops' conformational changes upon binding played a role. The comparison of conformational changes between bound (green) and unbound (blue) TCR and unbound pMHC (blue) prior to docking observed in X-ray experimental structures reveals significant structural shifts in specific loops and peptides across various protein complexes (**Suppl. Fig. S2**):

(A) Major conformational changes in CDR3α loops and peptides are observed in PDB ID 2NX5.
(B) Notable changes in CDR3β and CDR2β are seen in PDB IDs 4JFD and 4JFF.
(C) Alterations in CDR3α, CDR2β, and the peptide are present in PDB ID 6EQB.
(D) Shifts in CDR3α and peptide loops with long CDR3 loops (>=12 residues) are noted in PDB IDs 3KPR, 3KPR, and 1MI5.

These cases primarily produced acceptable-quality models rather than higher quality models (with the exception for 2NX5, which included medium-quality models in top 50 models) in both sampling and scoring (**Suppl. Fig. S1** and **Fig. 1**). In summary, two key factors may contribute to SwiftTCR generating fewer high-quality models in certain cases: (1) the presence of long CDR3 loops, and (2) inward-bending unbound CDR2 loops that create steric clashes with the MHC molecule.

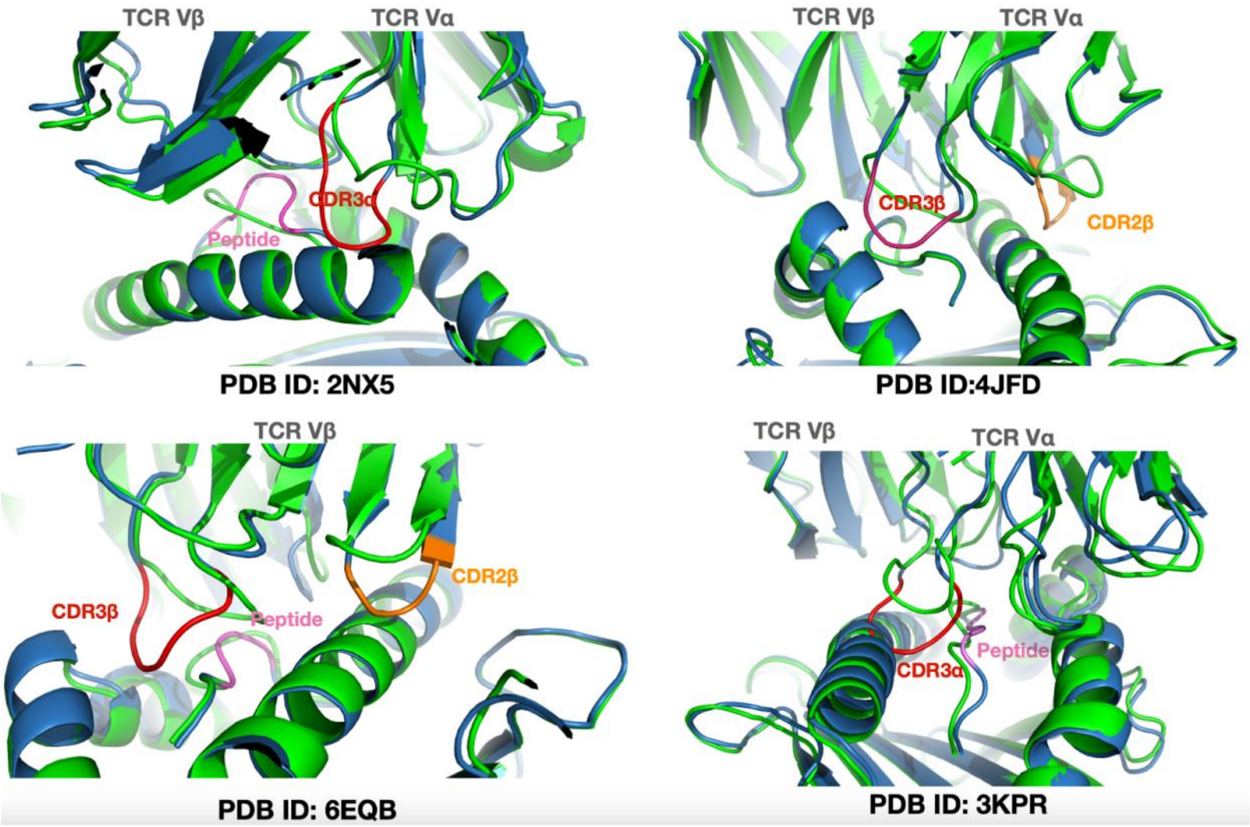


**Supplementary Figure S3. Comparison of conformational changes between bound (green) and unbound (blue) states observed in X-ray experimental structures.** A) Conformational changes in the CDR3α loops and the peptide in 2NX5. B) Conformational changes in CDR3β and CDR2β in 4JFD. C) Conformational changes in CDR3α, CDR2β, and the peptide in 6EQB. D) Conformational changes in long CDR3α and peptide loops in 3KPR.

#### 2.3 Electrostatic-favored energy weighting improves TCRpMHC docking accuracy

The strong negative correlation between CDR3 and peptide charges reported in previous work (Dash et al., 2017) suggests that electrostatics may be a key determinant in TCR–peptide recognition. We experimented with different weights for different energy terms, including van der Waals (vdW), electrostatics and pairwise potentials (**Suppl. Table 2**). **Suppl. Table 2** lists four different PIPER energy schemes with different weights. Ft 00 is the default PIPER energy function. Ft 02, Ft 03 and Ft 07 each have a higher weight assigned to electrostatics compared to Ft 00. We compared the three other configurations with the default weight set (Ft 00). The quality of the models generated by each configuration was evaluated to determine the best-performing set of weights.

Our results demonstrated improved performance with the electrostatic-favored energy function (**Fig. S3**). Specifically, Ft 02 showed a general enhancement in model quality, outperforming both the general energy function (Ft 00) and other weighting schemes (Ft 03 and Ft 07). While Ft 03 and Ft 07 also prioritized electrostatics, they assigned lower weights to attractive van der Waals energy and pairwise potential energy (**Suppl. Table 2**), leading to an overall reduction in model quality compared to Ft 00 (**Fig. S3**). Therefore, we set Ft 02 as the default energy scheme of SwiftTCR.

**Supplementary Table S3. Different PIPER energy weights.** The first column lists the weight names used in PIPER. The rest of the columns are weights for different energy terms. These weights are used to calculate the unweighted values, and the combination of all weighted values forms the total energy.

| Weight ID | Repulsive vdW energy | attractive vdW energy | coulombic electrostatic energy | Generalized born approximation electrostatics energy | pairwise potential energy |
|---|---|---|---|---|---|
| Ft 00 | 0.40 | -0.40 | 300 | 30 | 1.00 |
| Ft 02 | 0.40 | -0.40 | 600 | 60 | 1.00 |
| Ft 03 | 0.40 | -0.10 | 600 | 60 | 1.00 |
| Ft 07 | 0.40 | -0.10 | 600 | 60 | 0.20 |

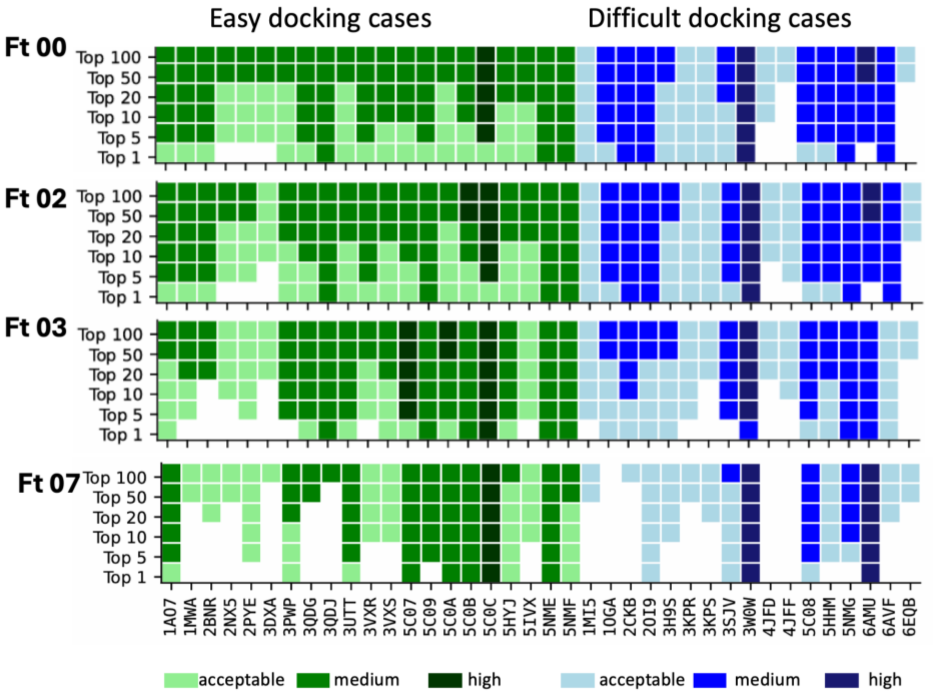


**Supplementary Figure S4. Comparison of different PIPER weights.** The results show that Ft 00 and Ft 02 perform similarly, while Ft 03 generally worsens compared to Ft 00 and Ft 07 performs the worst overall, frequently failing to generate acceptable-quality models.

#### References

Basu, S., & Wallner, B. (2016). DockQ: A Quality Measure for Protein-Protein Docking Models. *PLOS ONE*, *11*(8), e0161879. https://doi.org/10.1371/journal.pone.0161879

Dash, P., Fiore-Gartland, A. J., Hertz, T., Wang, G. C., Sharma, S., Souquette, A., Crawford, J. C., Clemens, E. B., Nguyen, T. H. O., Kedzierska, K., La Gruta, N. L., Bradley, P., & Thomas, P. G. (2017). Quantifiable predictive features define epitope-specific T cell receptor repertoires. *Nature*, *547*(7661), 89–93. https://doi.org/10.1038/nature22383

Dunbar, J., & Deane, C. M. (2016). ANARCI: Antigen receptor numbering and receptor classification. *Bioinformatics*, *32*(2), 298–300. https://doi.org/10.1093/bioinformatics/btv552

Kuffner, J. J. (2004). Effective sampling and distance metrics for 3D rigid body path planning. *IEEE International Conference on Robotics and Automation, 2004. Proceedings. ICRA '04. 2004*, 3993-3998 Vol.4. https://doi.org/10.1109/ROBOT.2004.1308895

Lensink, M. F., & Wodak, S. J. (2010). Docking and scoring protein interactions: CAPRI 2009. *Proteins: Structure, Function, and Bioinformatics*, *78*(15), 3073–3084. https://doi.org/10.1002/prot.22818

Lin, V., Cheung, M., Gowthaman, R., Eisenberg, M., Baker, B. M., & Pierce, B. G. (2025). TCR3d 2.0: Expanding the T cell receptor structure database with new structures, tools and interactions. *Nucleic Acids Research*, *53*(D1), D604–D608. https://doi.org/10.1093/nar/gkae840

Méndez, R., Leplae, R., Maria, L. D., & Wodak, S. J. (2003). Assessment of blind predictions of protein–protein interactions: Current status of docking methods. *Proteins: Structure, Function, and Bioinformatics*, *52*(1), 51–67. https://doi.org/10.1002/prot.10393

Rodrigues, J. P. G. L. M., Teixeira, J. M. C., Trellet, M., & Bonvin, A. M. J. J. (2018). pdb-tools: A swiss army knife for molecular structures. *F1000Research*, *7*, 1961. https://doi.org/10.12688/f1000research.17456.1

Shoemake, K. (1992). UNIFORM RANDOM ROTATIONS. In *Graphics Gems III (IBM Version)* (pp. 124–132). Elsevier. https://doi.org/10.1016/B978-0-08-050755-2.50036-1
